# Multi-omic subtypes of Alzheimer’s dementia are differentially associated with psychological traits

**DOI:** 10.1101/2025.02.21.639584

**Authors:** Andrea R. Zammit, Lei Yu, Victoria N. Poole, Konstantinos Arfanakis, Julie A. Schneider, Vladislav A. Petyuk, Philip L. De Jager, Rima Kaddurah-Daouk, Yasser Iturria-Medina, David A. Bennett

## Abstract

**Importance:** Psychological traits reflecting neuroticism, depressive symptoms, loneliness, and purpose in life are risk factors of AD dementia; however, the underlying biologic mechanisms of these associations remain largely unknown.

**Objective:** To examine whether one or more multi-omic brain molecular subtypes of AD is associated with neuroticism, depressive symptoms, loneliness, and/or purpose in life.

**Design:** Two cohort-based studies; Religious Orders Study (ROS) and Rush Memory and Aging Project (MAP), both ongoing longitudinal clinical pathological studies that began enrollment in 1994 and 1997.

**Setting:** Older priests, nuns, and brothers from across the U.S. (ROS) and older adults from across the greater Chicago metropolitan area (MAP).

**Participants:** 822 decedents with multi-omic data from the dorsolateral prefrontal cortex.

**Exposure(s):** Pseudotime, representing molecular distance from no cognitive impairment (NCI) to AD dementia, and three multi-omic brain molecular subtypes of AD dementia representing 3 omic pathways from no cognitive impairment (NCI) to AD dementia that differ by their omic constituents.

**Main outcome(s) and measure(s):** We first ran four separate linear regressions with neuroticism, depressive symptoms, loneliness, purpose in life as the outcomes, and pseudotime as the predictor, adjusting for age, sex and education. We then ran four separate analyses of covariance (ANCOVAs) with Bonferroni-corrected post-hoc tests to test whether the three multi-omic AD subtypes are differentially associated with the four traits, adjusting for the same covariates.

**Results:** Pseudotime was positively associated (*p*<0.05) with neuroticism and loneliness. AD subtypes were differentially associated with the traits: AD subtypes 1 and 3 were associated with neuroticism; AD subtype 2 with depressive symptoms; AD subtype 3 with loneliness, and AD subtype 2 with purpose in life.

**Conclusions and Relevance:** Three multi-omic brain molecular subtypes of AD dementia differentially share omic features with four psychological risk factors of AD dementia. Our data provide novel insights into the biology underlying well-established associations between psychological traits and AD dementia.

**Key points:** *Question:* Are three distinct multi-omic brain molecular subtypes of Alzheimer’s disease (AD) dementia associated with four well-established psychological AD risk factors (neuroticism, depressive symptoms, loneliness and purpose in life)?

*Findings:* We found differential associations: AD subtypes 1 and 3 were associated with neuroticism, AD subtype 2 was associated with depressive symptoms, AD subtype 3 was associated with loneliness; and AD subtype 2 was associated with purpose in life.

*Meaning:* Psychological risk factors might be associated with AD dementia via shared multi-omic molecular pathways.

## Introduction

Psychological traits, such as proneness to psychological distress (i.e., neuroticism), depressive symptoms, loneliness, and purpose in life, are well-established risk factors for prospective mental and physical health outcomes^1–5^ such as incidence of mild cognitive impairment (MCI) and Alzheimer’s disease and related dementias (AD/ADRD)^6–10^ as well as morbidity and mortality.^11–16^ The public health significance of these traits as robust risk factors has also been well-documented.^17–21^

We previously documented the associations of neuroticism^22^, depressive symptoms^23^, loneliness^24^, and purpose in life^25^, with cognitive decline, and incidence of MCI and AD/ADRD. These associations remained after adjusting for common age-related neuropathologic indices that cause cognitive impairment.^22–29^ To date we have not found any associations between these traits and common markers of neurodegeneration including neuritic plaques, neurofibrillary tangles, Lewy bodies, TDP-43, and hippocampal sclerosis, though we did find weak associations between purpose in life and lacunar cerebral infarctions.^30^ Thus, other molecular mechanisms are likely involved. For example, we reported that two neocortical proteins^31^ and a transcriptomic co-expression module^32^ mediated in part the association of neuroticism with cognition. Further, we also found some relationships linking depressive symptoms to cognition via select proteins^33^ and microRNAs.^34^

We recently integrated four layers of dorsolateral prefrontal cortex (DLPFC) omics data, i.e., epigenomic, transcriptomic, proteomic, and metabolomic, with multimodal contrastive Trajectories Inference (mcTI) analysis to derive multi-omic pseudotime from no cognitive impairment (NCI) to AD dementia^35^. We subsequently decomposed pseudotime into three distinct multi-omic brain molecular subtypes representing three molecular pathways from NCI to AD dementia. These subtypes differed in their multi-omic composition and key drivers. Here, we examined whether one or more of these molecular pathways would be associated with neuroticism, depressive symptoms, loneliness and/or purpose in life. Such associations would provide clues to potential molecular omic connections linking these traits to AD dementia.

## Methods

### Study population

Participants were community-based older adults from the Religious Orders Study (ROS) or the Rush Memory and Aging Project (MAP)^36^. ROS, initiated in 1994, includes older priests, nuns, and brothers from across the U.S. while MAP established in 1997, includes older men and women from across the greater Chicago metropolitan area. Participants were free of known dementia at enrollment, agreed to annual clinical evaluation and signed informed consent and Anatomic Gift Act to donate their brains at death. Both studies were approved by an Institutional Review Board of Rush University Medical Center.

### Assessment of psychological risk factors

Neuroticism was assessed using either 12 or 6 items from the NEO Five-Factor Inventory, which was administered at baseline or near baseline as previously reported^37^. Depressive symptoms were assessed annually using a 10-item form^38^ of the Center of Epidemiologic Study-Depression Scale(CES-D)^39^. Loneliness and purpose in life were assessed at baseline in MAP participants only, using a 5-item version from a modified scale of the de Jong-Gierveld Loneliness scale^40, 41^ and a 10-item scale derived from Ryff’s Scales of Psychological Wellbeing as previously described.^42, 43^ **eMethods** details psychometric information.

### Clinical Diagnoses

AD dementia was diagnosed by an experienced clinician using criteria of the joint working group of the National Institute of Neurologic and Communicative Disorders/Stroke/AD and Related Disorders Association^44^, as previously described.^45^ These criteria require a history of cognitive decline and evidence of impairment in at least two domains of cognitive function, one of which must be memory.^46^ MCI required evidence of impairment without meeting accepted criteria for dementia, and NCI refers to individuals without dementia or MCI as previously established. ^46, 47^

### Neuropathologic evaluation

Brain autopsy followed standardize protocols.^48, 49^ Neuropathologic evaluations systematically assessed common AD and non-AD neurodegenerative and cerebrovascular conditions including Alzheimer’s disease pathology, Lewy bodies, transactive response DNA binding protein (TDP)-43, hippocampal sclerosis, chronic macroscopic and microinfarcts, cerebral amyloid angiopathy(CAA), atherosclerosis, and arteriolosclerosis, as described in more detail in the **eMethods**.

### Postmortem Imaging

A subset of 278 participants underwent a postmortem brain imaging protocol, described previously^50^. Briefly, after one month postmortem, the cerebral hemisphere selected for neuropathological examination was imaged with a multi-echo spin-echo sequence on one of four 3-Tesla MRI scanners, and the resulting images were used for deformation-based morphometry(DBM) as detailed in **eMethods**.

### Annotation of the previously reported brain multi-omic molecular AD subtypes

Brain multi-omic data contain DNA methylation (DNAm) with Illumina 450 array, bulk next generation RNA sequencing, targeted protein expression with selected reaction monitoring, and metabolite level (metabolon) from DLPFC, generated as described previously. ^51–53^ We previously used the machine-learning mcTI algorithm to generate a brain molecular pseudotime of AD dementia relative to NCI, and three brain molecular subtypes reflecting different molecular pathways from NCI to AD dementia based on the numbers and types of omics.^35^ Here, we generate the top (FDR *p*<0.05) molecular features that characterize drivers of the three previously reported AD subtypes.^35^

### Statistical analyses

First, as pseudotime and the subtypes are built from AD dementia as the target, we compared frequency of pathologies across subtypes to ensure that our molecular subtypes were not simply capturing different pathological features. Pathologies were grouped accordingly:i) AD, CAA; ii)non-AD neurodegenerative: neocortical Lewy bodies, TPD-43, hippocampal sclerosis; iii) total infarcts, iv)cerebral vessel pathology: arteriosclerosis, atherosclerosis. We performed chi-squared tests to determine whether observed frequencies of pathologies differ by subtype.

Second, we described the proportion of the top epigenomic, transcriptomic, proteomic, and metabolomic features of the trajectory inferences per subtype, and illustrated their top 25 influential contributing features ordered by their F-value. Finally, we ran four separate analyses of variance tests, one for each omic modality (F-value as outcome and AD subtypes as predictors) using Bonferroni’s method for post-hoc analyses to calculate inter-subtype differences in the F-value.

Third, we tested associations with brain morphometry by running a general linear model of voxel-wise deformation assessed using FSL PALM^54, 55^ adjusting for demographics, postmortem interval, and scanner(accounting for differences in both mean and variance across scanners). We then contrasted each group pair. P-values were computed from 500 permutations using tail approximation^56^. The threshold-free cluster enhancement (TFCE) approach was used to define clusters of significance. Associations were considered statistically significant at p≤.05, family-wise error rate (FWER) corrected.

Fourth, we tested associations between pseudotime and the psychological traits in four separate linear regression models. Our outcomes were neuroticism, depressive symptoms, loneliness, and purpose in life. Covariates included age at death, sex, and education. The term for pseudotime indicated its association with the traits by one additional point.

Finally, we ran four analyses of covariance using Bonferroni’s method for multiple comparisons to test whether the 3 subtypes are associated with the psychological traits, adjusting for the same covariates.

## Results

A total of 822 participants had multi-omic data. Of those, 761 participants also had measures of neuroticism, 818 had CES-D, and 306 participants had measures on loneliness and purpose in life which were only available in MAP. Mean age at baseline was just over 80 years and mean age at death was close to 90 years. Over 60% were female, and most participants completed college(**Table 1**).

**Table 1.** Characteristics of the whole sample and stratified by controls and AD subtypes.

| Characteristics | Mean (SD) or % (n) |  |  |  |  |
| --- | --- | --- | --- | --- | --- |
|  | Whole sample | NCI | Subtype 1 | Subtype 2 | Subtype 3 |
| N | 822 | 274 | 189 | 197 | 162 |
| Age at baseline, mean years, (SD) | 82.3 (6.7) | 79.2 (6.9) | 81.9 (6.3) | 82.2 (6.8) | 82.9 (6.9) |
| Age at death, mean years (SD) | 88.4 (6.6) | 86.2 (6.5) | 89.3 (6.4) | 89.1 (6.6) | 90.4 (6.1) |
| Female, n (%) | 532 (64) | 171 (32) | 130 (24) | 122 (22) | 109 (20) |
| Educational attainment, mean years (SD) | 16.3 (3.5) | 16.3 (3.6) | 16.4 (3.6) | 16.2(3.5) | 16.3 (3.2) |
| Neuroticism, baseline, (SD) | 16.8 (6.6) | 15.76 (6.68) | 17.83 (6.61) | 16.72 (6.39) | 17.53 (6.26) |
| Depressive symptoms, mean score (SD) | 1.44 (1.37) | 1.26 (1.26) | 1.52 (1.44) | 1.58 (1.37) | 1.48 (1.46) |
| Loneliness, baseline (SD) | 2.4 (0.6) | 2.2 (0.6) | 2.4 (0.6) | 2.3 (0.5) | 2.5 (0.7) |
| Purpose in life, baseline (SD) | 3.6 (0.5) | 3.6 (0.5) | 3.5 (0.4) | 3.5 (0.4) | 3.5 (0.4) |
| Pseudotime, mean (SD) | 0.4 (0.21) | 0.23 (0.13) | 0.38 (0.13) | 0.45 (0.09) | 0.68 (0.18) |
| Global cognition, baseline, mean (SD) | -0.23 (0.7) | 0.17 (0.4) | -0.40 (0.6) | -0.45 (0.7) | -0.43 (0.9) |
| Global cognition, last valid, mean (SD) | -0.96 (1.2) | 0.08 (0.4) | -1.45 (1.1) | -1.50 (1.1) | -1.50 (1.1) |
| Mild cognitive impairment proximate to death, n (%) | 197 (23) | 0 | 63 (31) | 78 (39) | 56 (28) |
| AD dementia proximate to death, n (%) | 351 (42) | 0 | 126 (35) | 119 (33) | 106 (30) |
| Neuropathology |  |  |  |  |  |
| Pathologic AD (NIA-AA), n (%) | 517 (62.9) | 118 (43.1) | 141 (74.6) | 138 (70.1) | 120 (74.1) |
| Neocortical Lewy bodies, n (%) | 84 (10.2) | 13 (4.7) | 18 (9.5) | 28 (14.2) | 25 (15.4) |
| TDP-43, n (%) | 238 (30.4) | 43 (16.5) | 64 (36.4) | 72 (38.3) | 59 (37.3) |
| Hippocampal sclerosis, n (%) | 60 (7.4) | 6 (2.2) | 17 (9.0) | 25 (12.8) | 12 (7.5) |
| Gross chronic infarcts, n (%) | 294 (35.8) | 61 (22.3) | 83 (43.9) | 77 (39.1) | 73 (45.1) |
| Micro chronic infarcts, n (%) | 225 (27.4) | 63 (23.0) | 60 (31.7) | 56 (28.4) | 46 (28.4) |
| Arteriosclerosis, moderate to severe, n (%) | 317 (38.9) | 78 (28.7) | 74 (39.4) | 99 (50.8) | 66 (41.3) |
| Atherosclerosis, moderate to severe, n (%) | 346 (42.2) | 86 (31.7) | 84 (44.4) | 98 (49.7) | 78 (48.1) |
| Cerebral amyloid angiopathy, moderate, n (%) | 281 (35.1) | 75 (28.2) | 69 (37.1) | 73 (38.4) | 64 (40.3) |

### Frequency of AD/ADRD pathologies across AD subtypes

The reference NCI group had less pathology, as expected (median=2 pathologies, interquartile range (IQR)=2). AD subtypes had a median of 3 pathologies (IQRs:subtype 1=2; subtypes 2/3=3). Over half of NCI participants had 1 or 2 pathologies, while a third had three or more. Meanwhile over two thirds of participants in all AD subtypes had three or more pathologies, which is expected since mixed pathologies commonly drive AD dementia**(eTable 1).**

Over 70% of individuals across all subtypes had pathologic AD and/or CAA (χ^2^(2, 535)=1.96, *p*=0.38), and almost half had non-AD neurodegenerative neuropathology (χ^2^(2, 517)=4.09, *p*=0.66). There were no differences in the distribution of infarcts ((χ^2^(2, 548)=4.01,*p*=0.14) or in frequency of cerebral vessel neuropathology (χ^2^(4, 543)=2.89,*p*=0.24)(**Figure 1**).Therefore, our molecular subtypes are not simply capturing different neuropathological features.

**Figure 1.**
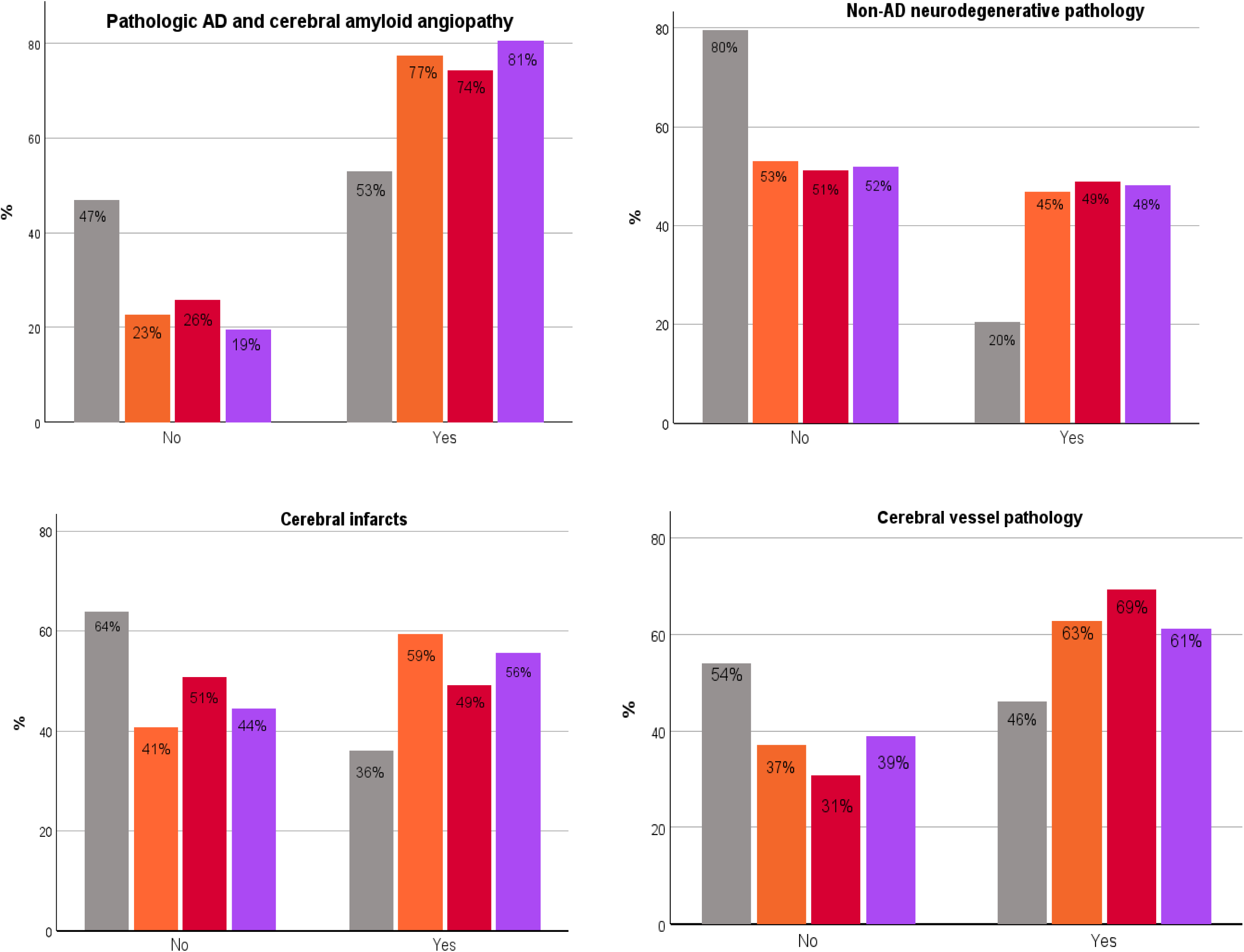
Percentage of individuals with ADRD pathologic indices stratified by No cognitive impairment and multi-omic brain molecular AD subtypes. Pathologic indices include diagnosis of AD, non-AD neurodegenerative pathology (neocortical Lewy bodies, TPD-43, and hippocampal sclerosis), cerebral (gross and/or micro) infarcts, and cerebral vessel pathology (arteriosclerosis, atherosclerosis, and cerebral amyloid angiopathy). Different colored bars represent the percentage of individuals within the no cognitive impairment (NCI) group and each AD subtype and add up to 100% for each individual group. *Note.* Gray = no cognitive impairment; Orange= AD subtype 1; Red=AD subtype 2; Purple=AD subtype 3.

### Proportion of omic features for each AD subtype

The relative contribution of each omic dimension was related to abundance with the greatest number of features: DNAm, followed by bulk RNAseq, and targeted proteomics which had the fewest. Thus, most frequent omic features across all subtypes were epigenomic alterations, followed by RNA alterations which is expected as they are the two with by far the greatest number of features. Still, there were significant differences in omic proportions amongst the AD subtypes (χ^2^(6,569)=30.4, *p*<0.001)(**Figure 2a**). For example, while 15% of the top features in subtype 1 were metabolites, less than 5% in subtypes 2 and 3 were metabolites. Almost 43% of top features in subtype 2 were RNA alterations while in subtypes 1 and 3 these alterations amounted to about 30%. Finally, while over 60% of epigenomic alterations characterized subtype 3, these were less then 50% in subtypes 1 and 2. Proteomic abnormalities contributed the least, as expected given the small number of features.

**Figure 2.**
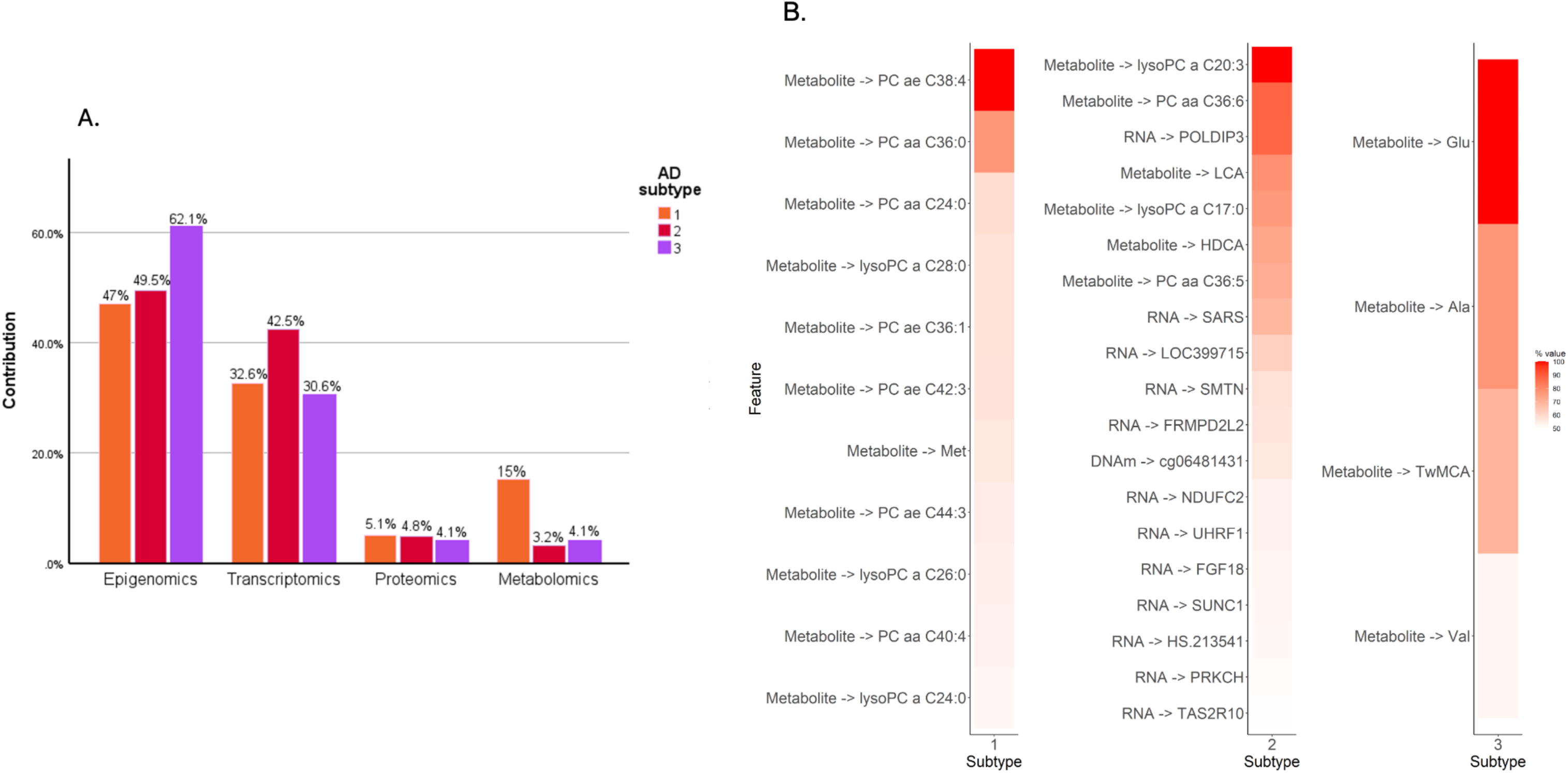
Modality-specific omic contributions and top contributing omic markers per AD subtype. **Panel A. Modality-specific contributions (in percentages) of epigenomic, transcriptomic, proteomic, and metabolomic alterations relative to the three identified AD subtypes. Each colored bar adds up to 100% for that particular AD subtype.** Therefore, while epigenomic features were the most common omic modality in all subtypes, relative to each other, subtype 1 was characterized by metabolomic alterations, subtype 2 by transcriptomic alterations, and subtype 3 by epigenomic alterations. Only markers significantly associated at FDR *p* < 0.05 were considered in this study. **Panel B. Top molecular omics contributions to AD molecular subtypes in the DLPFC of the postmortem human brain.** Top influential metabolomic, transcriptomic, epigenomic and proteomic markers per AD subtype (F-values in percentages, only F-values>50% of omic signals are shown). In all, while epigenomic features were the most frequent data modality across subtypes, the strongest contributors as determined by the F-value, consistently were metabolites.

### Top omic features for each AD subtype

Top influential contributing omic features with an F-value>50% can be seen in **Figure 2b**. Interestingly, almost all the strong drivers were metabolites. The strongest contributors (F-value>90%) differentiating these AD subtypes from NCI were three phospholipids and a major excitatory neurotransmitter. Specifically, phosphatidylcholine acyl-alkyl (PCae)C38:4 for AD subtype 1; lysophosphatidylcholine (lysoPC)acyl C20:3 and PC acyl-acyl (aa)C36:6 for AD subtype 2; and glutamate for AD subtype 3 (full list in **eTable 2)**.

### Subtype-subtype differences of top influential omic features by modality

We then compared each omic modality’s F-values amongst subtypes; higher F-values indicate stronger differences from NCI relative to other subtypes. AD subtype 2 consistently had significantly higher F-values across all omic features (**Figure 3**). Specifically, epigenomic alterations were significantly higher in AD subtype 2 relative to subtypes 1 and 3 as were RNA alterations, proteomic alterations, and metabolomic alterations. **eTable 3** lists means and 95%CIs. In sum, the top omic features in subtype 2 differentiated this subtype from NCI more strongly than either of the omic features in subtypes 1 and 3. Yet, specific omic features at the individual level were different per subtype. **eTable 4** details mean differences.

**Figure 3.**
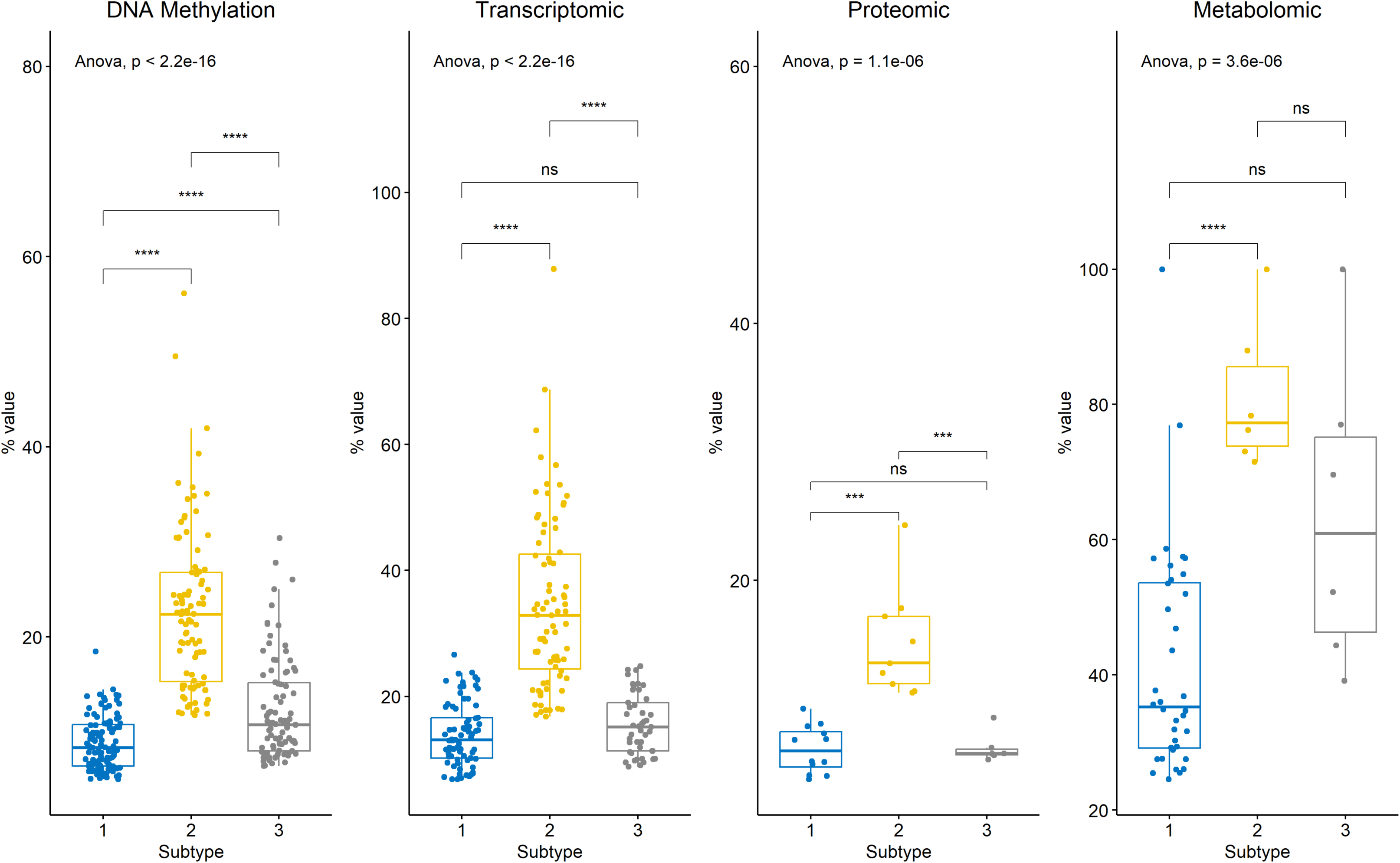
Box and whisker plots of inter-subtype molecular differences. Each omic data feature shows a comparison amongst the three AD subtypes. Each plot corresponds to the percent of significantly different features in percent F-value for each omic modality, based on ANOVA tests with subtype as the grouping measure. The dots in each plot represent a feature within that omic modality; the top and bottom of the boxes represent quartiles 1 and 3, and the whiskers show the inter-quartile range. The middle horizontal line represents the mean percent of data features that are abnormal for that subtype.

### Brain morphology across AD subtypes

To further characterize the subtypes, we examined differences in brain morphometry across a subset of 278 participants (NCI=91, AD subtypes: 1 n=74; 2 n=57, 3 n=56) with postmortem MRI. As expected, all AD subtypes exhibited greater cortical atrophy, as indicated by smaller postmortem brain volumes, relative to NCI (**eFigure 1**). AD subtype 1 had the most extensive atrophy, as observed by smaller volumes in the medial temporal lobe and across several other temporal, frontal, and parietal regions. AD subtype 2 exhibited temporal lobe atrophy that extended to the temporal pole. Finally, AD subtype 3 exhibited the least atrophy, which was largely localized to the temporal lobe and insular cortex. Both subtypes 1 and 2 also had extensive ventricular enlargement, which indicated central brain atrophy, while subtype 3 had minimal enlargement of the temporal horn. While the differences are intriguing, none of the subtypes were statistically different from one another in this small subsample.

### Association of pseudotime with psychological traits

Mean pseudotime in the NCIs was almost 0.25 while across the AD subtypes this ranged from about 0.4 for subtype 1 to 0.5 for subtype 2 and 0.7 for subtype 3, with higher mean pseudotime indicating closer proximity to AD dementia(**Table 1**).

Higher pseudotime was positively associated with neuroticism (beta=2.86, 95%CI=0.62-5.10, *p*=0.013) and loneliness (beta=0.35, 95%CI=0.04-0.66, *p*=0.029). Pseudotime explained 4% of variance in neuroticism, and 3% of variance in reported loneliness after accounting for age, sex, and level of education. There was no association between pseudotime and depressive symptoms (beta*=*0.402, 95%CI = −0.052–0.856, *p*=0.082), or purpose in life(beta=-0.15, 95%CI=-0.38-0.08, *p*=0.212).

### Association of AD subtypes with psychological traits

ANCOVA showed significant effects of the subtypes on all traits (neuroticism: F(3,760)=5.9, *p*<0.001; depressive symptoms: F(3,817)=3.9, *p*=0.009; loneliness: F(3,305)=5.6, *p*=0.014; purpose in life, F(3,305)=3.5, *p*=0.016).

Post-hoc comparisons indicated differential associations between AD subtypes and the traits relative to NCI; however, there were no inter-subtype differences(**Figure 4**). Subtypes 1 and 3 had higher neuroticism scores than NCI, while subtype 2 had higher depressive symptoms. Subtype 3 had higher loneliness; subtype 2 had marginally lower purpose in life score(**eTable 5**).

**Figure 4.**
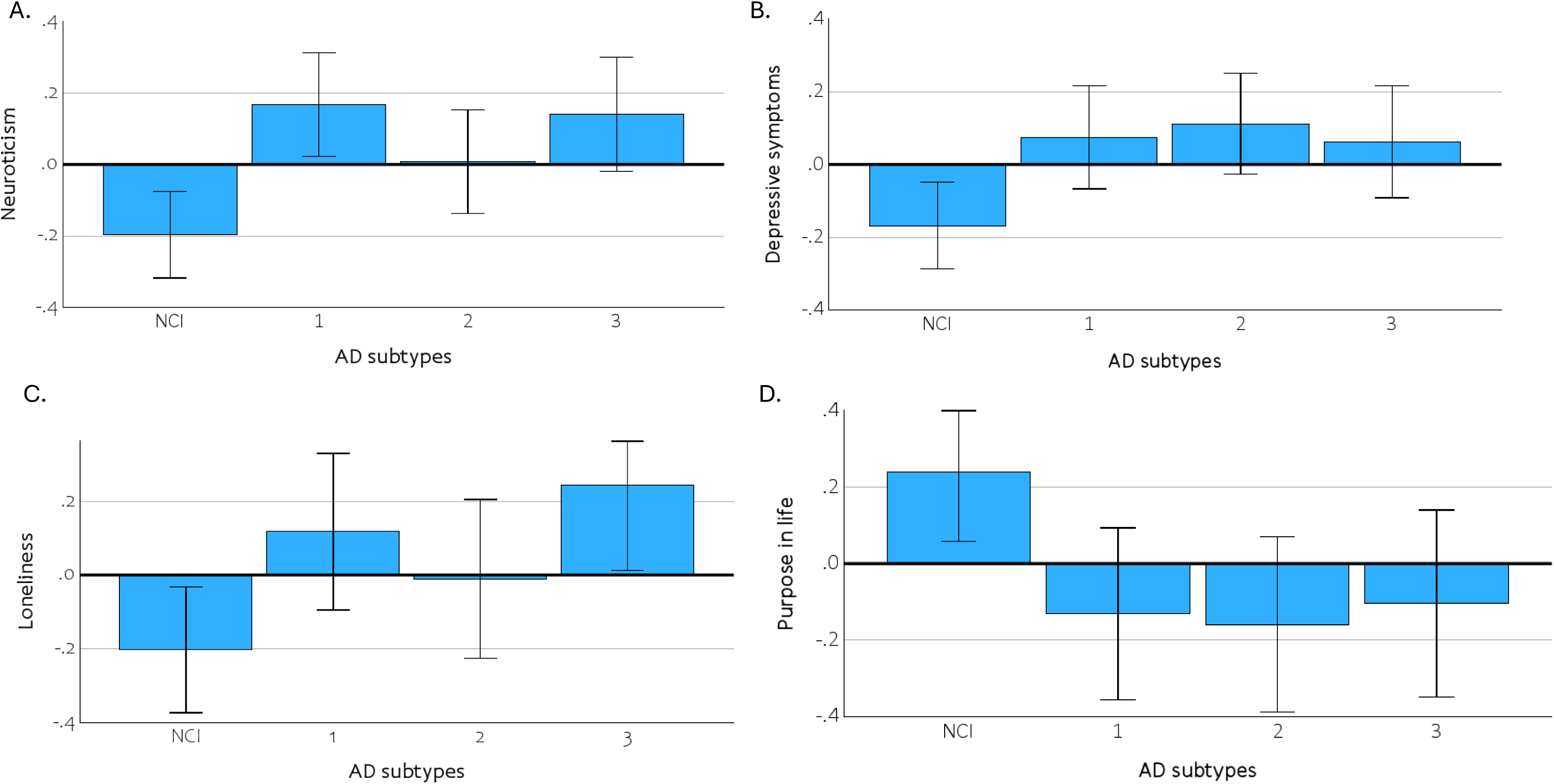
Bar charts illustrating distance from the mean of each psychological risk factor (panels A-D). Results are stratified by the no cognitive impairment (NCI) group and the three AD subtypes. The measures are z-transformed for illustration purposes. The means are adjusted for age, sex, and education.

## Discussion

To better understand the mechanistic basis of the relation of four psychological traits to AD dementia, we leveraged prior work in 822 older adults who subsequently died and had brain autopsy, providing brain multi-omic molecular pseudotime, and three distinct molecular subtypes representing pathways from NCI to AD dementia. We found that pseudotime was associated with neuroticism and loneliness, and subsequently AD multi-omic brain molecular subtypes were differentially associated with neuroticism, depressive symptoms, loneliness, and purpose in life. The results provide novel data on shared top multi-omic features between molecular subtypes of AD dementia and well-established AD/ADRD risk factors, suggesting common underlying mechanisms driving associations between psychological traits and AD/ADRD.

Previous studies primarily focused on single-omic associations. We previously showed that neuroticism is associated with 18 cortical transcriptomic co-expressed modules, three (m6, m7, and m127) of which partially mediated the association of higher neuroticism on faster cognitive decline^32^. We previously also identified two cortical proteins, 40S ribosomal protein S3 and BCKDHB, that were associated with both neuroticism and cognitive decline independent of common AD/ADRD pathologic indices.^31^ These two proteins strongly mediated the association of higher neuroticism on cognitive decline independent of common AD/ADRD pathologic indices, and further explained 25% of the variance in this association. Together, our prior studies provide evidence that neuroticism has strong and widespread effects on the transcriptome, and on specific cortical proteins in the aged prefrontal cortex, both of which affect downstream AD-related outcomes. Similarly, in prior work we identified a cortical protein, IGFBP-5, that was associated with both depressive symptoms and cognition, and further explained 10% of the association between depressive symptoms and cognitive decline.^33^ We also previously identified four microRNAs associated with depressive symptoms, of which miR-484 targets were enriched in a co-expression module involved in synaptic function and plasticity and were associated with higher risk of AD dementia.^34^ While there is some evidence on associations of loneliness^57^, and purpose in life^58^, with altered DNAm, and expression of mRNAs and proteins, their shared mechanisms with AD/ADRD are still largely unexplored.

We extend prior work by showing here that including multiple molecular modalities to identify common correlates of subtypes representing distinct molecular signatures of AD dementia can be leveraged to potentially identify the molecular basis of psychological traits associated with AD dementia. To our knowledge, this is the first study that has leveraged multi-omics data to directly explore these associations. This approach allowed us to identify distinct molecular AD subtypes that are differentially associated with four psychological traits. AD subtypes 1 and 3 showed strong associations with neuroticism, while subtype 2 was exclusively associated with depressive symptoms. AD subtype 1 had higher proportions of metabolomic alterations relative to subtypes 2 and 3, while subtype 3 had higher proportions of epigenomic alterations. By contrast, subtype 2 had relatively higher proportions of transcriptomic alterations. Loneliness and purpose in life had somewhat weaker associations with subtypes 3 and 2 respectively likely due to low statistical power since these two measures were only available in MAP participants. Much work remains to expand our findings in better-powered studies, with additional and larger multi-layered omic subtypes.

Our findings suggest that the various AD molecular mechanisms that are subserving cognition are also related to psychological traits. Metabolomic alterations were the topmost features that significantly differentiated these AD subtypes from NCI. Top alterations in subtypes 1 and 2 were phosphatidylcholines (PCs), a major class of phospholipids that form a crucial component of cell membranes. PC acyl-acyl (AA) and acyl-ether (AE) lipids are involved in signaling and maintenance of membrane structure; the ether bond in the AE lipids further promote resistance to oxidative damage. Disruption in lipid metabolism, including significant reductions in PC ae c38:4 and PC aa C36:6 are significantly associated with clinical AD.^59^ ^60, 61^ LysoPCs are proinflammatory lipids and their concentration is substantially higher in clinical AD.^62^ Psychological distress has also been associated with altered lipid metabolism via a cascade of dysregulation in PCs and lysoPCs that eventually cause cell membrane destabilization and inflammatory responses.^63^ The top alteration in AD subtype 3 was glutamate which is the primary excitatory neurotransmitter in the central nervous system. Overactivation of glutamate receptors are implicated in neuronal damage and death via a cascade of intracellular events including oxidative stress, mitochondrial dysfunction, and activation of destructive enzymes.^64^ Chronic distress and depression have also been implicated in the malfunctioning of the glutamatergic neurotransmitter system primarily affecting glutamate receptor expression in the prefrontal cortex, hippocampus and amygdala^65, 66^. Further studies that will collect more omics data will be better able to characterize multi-level omic subtypes and determine mediating effects of top contributors of AD subtypes on the associations between enduring psychological traits and AD/ADRD outcomes.

This study has limitations and strengths. The brief neuroticism measure used here may have underestimated associations, though we have previously reported a correlation of .90 between shorter-(6-and 12-) item versions of the NEO^22^ testifying to its validity. Nonetheless associations might be slightly stronger than what we observed here. Similarly, the depressive symptoms measure is also based on a brief psychometric measure, however, its reliability has been previously established^67^. For CES-D we used each person’s average score across all evaluations as previously^68^, which capture stable enduring tendencies, and reduce random error. Measures of loneliness and purpose in life were only available in MAP participants, potentially reducing statistical power. All omics came from one brain region and the number of epigenomic features and transcriptomic were disproportionally higher than those for metabolites or proteins^35^, which might have overestimated the noted proportions of epigenomic features within subtypes. However, the DLPFC is a known region for its associations with learning, stress response and emotion regulation, and the continuous collection of omics data within our cohorts will allow for further multilevel characterization of complex traits and replication of results. Our novel multi-omic approach provides insights into the molecular basis of psychological AD/ADRD risk factors. Other approaches have been used to identify molecular subtypes of AD; however, they have been agnostic to any clinical trait.^69–72^ High participation rates in the clinical evaluations and brain autopsy in the ROS and MAP cohorts minimize bias due to selective attrition.

## Data sharing

The data are available via the Rush Alzheimer’s Disease Center Research Resource Sharing Hub. Qualified applicants should complete an application including study premises and a brief description of the research plan.

Brain multi-omic data are available in the AMP-AD knowledge portal (www.synapse.org), using the following Synapse IDs:syn3157275 (epigenomic data), syn3800853 (transcriptomic), syn10468856 (proteomic), and syn10235595 and syn10235594 (metabolomic).

## Supporting information

eMethods

## Acknowledgements

We appreciate the participants of MAP for the time generously given for data collection and for consenting for brain donation. We also acknowledge staff of Rush Alzheimer’s Disease Center for data collection, management, and analyses.

This work was supported by National Institutes of Health: R01AG17917, P30AG10161, P30AG72975, R01AG015819, U01AG61356, U01 NS100599, R01 AG064233, RF1 NS139975, P30 AG072975. We would also like to thank Michael Urbut and the Paul M. Angell Family Foundation for their generous financial support given towards this work.

The funding organizations had no role in the design or conduct of the study; collection, management, analysis, or interpretation of the data; or preparation, review, or approval of the manuscript.

## Contributors

ARZ, YIT and DAB were responsible for the conception and design of the current study. ARZ performed the literature review and wrote the first draft of the manuscript. LY conducted the data analysis and contributed to writing the statistical analysis section and revising the manuscript. LY supervised the data analysis and contributed to revising the manuscript. KA is the director of the neuroimaging core of Rush Alzheimer’s Disease Center. KA and VNP supervised and oversaw the neuroimaging analyses. JAS is the director of pathology core of Rush Alzheimer’s Disease Center that collects indices of brain pathologies. She is also the director of Rush Alzheimer’s Disease Research Center that oversees data collection of the ROS cohort. VAP selected the protein targets and designed the peptides. PLD performed LC-SRM targeted proteomic assays and provided quantitative datasets of the peptide. RKD leads Alzheimer Disease Metabolomics Consortium and Alzheimer Gut Microbiome Project funded by NIA that contributed metabolomics data generated by Metabolon Inc. where metabolomics was conducted by chromatography spectrometry. DAB is the principal investigator of MAP and was involved in data collection. LY, VNP, KA, JAS, VAP, PLD, YIT and DAB contributed to revising the manuscript. YIT and DAB had full access to all the data in the study, accessed and verified the data, and take responsibility for the integrity of the data and the accuracy of the data analysis. All authors had access to the data and had final responsibility for the decision to submit for publication.

