## Supplementary material for "Multi-omic subtypes of Alzheimer’s dementia are differentially associated with psychological traits": eMethods

Both cohorts have identical study design and a large common core of data collection at the item level. The same trainer, supervisor and testers collect the data, and the brains go through the same neuropathologic and omic pipelines.

1. **Psychometric characteristics of the psychological measures**

**Neuroticism.** Neuroticism is an indicator of proneness to psychological distress. Participants rated agreement with each item (e.g. “I often feel inferior to others”), on a 5-point Likert scale ranging from 0 (strongly disagree) to 4 (strongly agree). The total score in participants with the 12-item scale (n=677) was computed by summing item scores to yield a total score that ranged from 0 to 48, with higher scores indicating more proneness to psychological distress. The total score of those with the 6-item version (n=84) was computed by adding the item scores and then multiplying the sum by two to make the score comparable to the standard 12-item version. The correlation of the 6-item version with the standard 12-item scale is 0.90, as previously reported^22^, supporting the validity of the 6-item scale. Internal consistency reliability as determined by the Cronbach coefficient alpha was 0.8, which indicates adequate internal consistency.

**Depressive symptoms.** The CES-D is widely used in epidemiologic studies of older persons, and the reliability of this version with its original version has been previously reported^38^. Each item represented a symptom, and participants indicated whether they experienced the symptom during the past week. The score was the total number of symptoms experienced. Because we did not observe any systematic change in CES-D scores in participants during follow-up^40^, we used each participant’s average CES-D score across all evaluations to enhance our assessment of the enduring disposition to experience depressive symptoms, while reducing measurement error and the impact of episodic memory fluctuations.

**Loneliness.** Participants rated agreement with each item (e.g. “I experience a general sense of loneliness.”) on a 5-point Likert scale ranging from 1 (strongly disagree) to 5 (strongly agree). These five items were selected to explicitly measure the psychological and feelings of isolation. The original version of this scale has been shown to be internally consistent^42, 43^.

**Purpose in life.** Participants rated agreement with each item (e.g. “I enjoy making plans for the future and working them to a reality.”) on a 5-point Likert scale ranging from 1 (strongly disagree) to 5 (strongly agree). The total score was then averaged to yield a total score, with higher scores indicating higher purpose in life.

2. **Neuropathologic evaluation**

Brain autopsy procedures including brain removal, tissue sectioning and preservation, and a uniform gross and microscopic examination with quantification of postmortem indices, have been described.^48, 49^ Briefly, brains were removed and weighed, and hemispheres were cut coronally using a Plexiglas^®^ jig into 1cm slabs. One hemisphere was fixed in 4% paraformaldehyde. After gross examination of both hemispheres, nine brain regions of interest (i.e. mid-frontal, mid-temporal, inferior parietal, anterior cingulate, entorhinal and hippocampal cortices, basal ganglia, thalamus, and mid-brain) were dissected from the 1cm slabs of fixed tissue and processed and embedded in paraffin. Sections (6mm) from the paraffin blocks were stained for assessment of pathology.

**AD and non-AD neurodegenerative pathologies.** For AD pathological diagnosis, a modified Bielschowsky silver stain was used to visualize neuritic plaques, diffuse plaques, and neurofibrillary tangles in five cortical areas (hippocampus, entorhinal, midfrontal, middle temporal, and inferior parietal). A board-certified neuropathologist or an expert neuroscientist, blinded to clinical data, determined the pathologic diagnosis of AD, defined as intermediate to high likelihood of AD according to NIA Reagan criteria^51^. The presence of neocortical Lewy body pathology was determined using antibodies to a-synuclein^52^. The presence of hippocampal sclerosis was determined using H&E stain^53^. The presence of TDP-43 cytoplasmic inclusions was determined using antibodies to phosphorylated TDP-43; TDP-43 distribution was grouped into 3 stages (stage 1, localized to amygdala; stage 2, extension to hippocampus or entorhinal cortex; stage 3, extension to the neocortex) and TDP-43 was considered present if positive for stages 2 or 3^54^.

**Vascular pathologies.** Gross chronic infarcts were identified by visually examining slabs and pictures from both brain hemispheres and confirmed histologically, and microinfarcts (chronic) were identified under microscopy using hematoxylin and eosin stain^49^. Moderate to severe arteriolosclerosis was detected by use of hematoxylin and eosin stained section of the anterior basal ganglia^55^. Moderate to severe atherosclerosis was detected by visual inspection of vessels in the Circle of Wilis^55^. Moderate to severe cerebral amyloid angiopathy was detected using amyloid-B immunostaining in four cortical regions^55, 56^.

3. **Postmortem Imaging**

The parameters of the ex-vivo MRI protocol were similar for the four MRI scanners used in this work, and have been presented in detail in reference 50. The ex-vivo brain MRI images were masked, corrected for field inhomogeneity using the N4 approach^51^, and were non-linearly registered to an in-house postmortem hemispheric template^52^ using Advanced Normalization Tools (ANTs)^53^. The determinant of the Jacobian of the transformation was calculated voxel-wise, and the resulting maps were smoothed with a 4mm FWHM Gaussian kernel. Higher values in these smoothed maps indicated the presence of more tissue that had to be contracted through image registration to fit the template, whereas lower values indicated less tissue that had to be expanded to fit the template.
