## Supplementary Material for "Multi-omic subtypes of Alzheimer’s dementia are differentially associated with psychological traits"

**Tables**

**eTable1.** Number of pathologies stratified by no cognitive impairment (NCI) and AD subtypes.

| Frequency of pathologies | NCI | AD subtype 1 | AD subtype 2 | AD subtype 3 |
| --- | --- | --- | --- | --- |
|  | N (%) | N (%) | N (%) | N (%) |
| 0 | 34 (13.4) | 1 (0.6) | 6 (3.3) | 6 (3.3) |
| 1 | 65 (25.7) | 25 (14.5) | 19 (10.6) | 19 (12.4) |
| 2 | 70 (27.7) | 34 (19.7) | 32 (17.8) | 27 (17.6) |
| 3 | 48 (19.0) | 37 (21.4) | 36 (20.0) | 30 (19.6) |
| 4 | 21 (8.3) | 36 (20.8) | 39 (21.7) | 33 (21.6) |
| 5 | 13 (5.1) | 28 (16.2) | 24 (13.3) | 27 (17.6) |
| 6 | 1 (0.4) | 6 (3.5) | 16 (8.9) | 10 (6.5) |
| 7 | 1 (0.4) | 6 (3.5) | 7 (3.9) | 2 (1.3) |
| 8 | 0 | 0 | 0 | 0 |
| 9 | 0 | 0 | 1 (0.6) | 0 |

**eTable 2.** Full list of omic signals contributing to the three AD subtypes at FDR *p* <0.05.

| **AD subtype 1** | **AD subtype 2** | **AD subtype 3** |
| --- | --- | --- |
| Metabolite -> PC ae C38:4; Metabolite -> PC aa C36:0; Metabolite -> PC aa C24:0; Metabolite -> lysoPC a C28:0; Metabolite -> PC ae C36:1; Metabolite -> PC ae C42:3; Metabolite -> Met; Metabolite -> PC ae C44:3; Metabolite -> lysoPC a C26:0; Metabolite -> PC aa C40:4; Metabolite -> lysoPC a C24:0; Metabolite -> PC aa C42:5; Metabolite -> PC ae C40:4; Metabolite -> PC ae C40:5; Metabolite -> Cit; Metabolite -> PC aa C26:0; Metabolite -> PC ae C36:0; Metabolite -> PC ae C38:3; Metabolite -> C14:2-OH; Metabolite -> C14:1-OH; Metabolite -> PC aa C42:4; Metabolite -> His; Metabolite -> PC ae C44:6; Metabolite -> PC ae C30:0; Metabolite -> PC ae C38:2; Metabolite -> C10; Metabolite -> PC ae C34:1; Metabolite -> PC aa C30:0; Metabolite -> C14:2; Metabolite -> lysoPC a C26:1; Metabolite -> Orn; Metabolite -> SDMA; Metabolite -> PC aa C40:3; Metabolite -> lysoPC a C28:1; Metabolite -> PC ae C36:4; Metabolite -> PC ae C36:2 (RNA -> FXYD7; RNA -> S100PBP; RNA -> LOC642450; RNA -> TFPT; RNA -> TRIML2; RNA -> C21ORF34; RNA -> GRK4; RNA -> WDR19; RNA -> PLAC8L1; RNA -> NSUN5C; RNA -> SNRP70; RNA -> MAST1; RNA -> TMEM50A; RNA -> FLJ39061; RNA -> MYADM; RNA -> FNBP4; RNA -> HS.391327; RNA -> MESP1; RNA -> HS.538763; RNA -> LOC651125; RNA -> HSPC111; RNA -> IDUA; RNA -> C21ORF114; RNA -> MGC27345; RNA -> LOC650155; RNA -> TNFRSF25; RNA -> LOC388237; RNA -> HS.458154; RNA -> HS.532239; RNA -> NPIP; RNA -> LST-3TM12; RNA -> PDE5A; RNA -> KRT13; RNA -> LAMA5; RNA -> LAMA3; RNA -> LOC731042; RNA -> ELL3; RNA -> NXF1; RNA -> TRAPPC6A; RNA -> PSMD5; RNA -> HS.576530; RNA -> CAPNS1; RNA -> CHKB; RNA -> HS.581313; RNA -> ABL2; RNA -> TAZ; RNA -> GOLGA8G; RNA -> PBX4; RNA -> SIL1; RNA -> HIST1H2BO; RNA -> MYOC; RNA -> HS.407028; RNA -> HSF4; RNA -> LOC647371; RNA -> ZNF649; RNA -> GTPBP8; RNA -> SCPEP1; RNA -> HS.105791; RNA -> LOC650887; RNA -> LOC652168; RNA -> POLR3F; RNA -> SETD2; RNA -> HS.570762; RNA -> ACADVL; RNA -> LOC644143; RNA -> TPPP3; RNA -> LOC642980; RNA -> TIMM17B; RNA -> TENC1; RNA -> C14ORF148; RNA -> MBTPS1; RNA -> PTPRE; RNA -> CCDC43; RNA -> FLJ41481; RNA -> GTF2IRD2B; RNA -> HS.559929; RNA -> HS.314177 (DNAm -> cg11278506; DNAm -> cg10104252; DNAm -> cg14074486; DNAm -> cg22176017; DNAm -> cg11995069; DNAm -> cg26709695; DNAm -> cg15850287; DNAm -> cg16668548; DNAm -> cg25664220; DNAm -> cg08277100; DNAm -> cg19360330; DNAm -> cg20911707; DNAm -> cg13027384; DNAm -> cg15077558; DNAm -> cg18574476; DNAm -> cg07285896; DNAm -> cg17204018; DNAm -> cg23517743; DNAm -> cg05005586; DNAm -> cg14664242; DNAm -> cg00678938; DNAm -> cg08005122; DNAm -> cg25061682; DNAm -> cg04134031; DNAm -> cg23894539; DNAm -> cg05809481; DNAm -> cg14270725; DNAm -> cg16652332; DNAm -> cg00844791; DNAm -> cg13638257; DNAm -> cg06002947; DNAm -> cg01809217; DNAm -> cg23002008; DNAm -> cg12833207; DNAm -> cg06565615; DNAm -> cg08030134; DNAm -> cg19131647; DNAm -> cg17696847; DNAm -> cg08787039; DNAm -> cg00167958; DNAm -> cg02186298; DNAm -> cg06837765; DNAm -> cg22425711; DNAm -> cg26875637; DNAm -> cg15964611; DNAm -> cg05682952; DNAm -> cg21951594; DNAm -> cg20801456; DNAm -> cg01233948; DNAm -> cg22222413; DNAm -> cg17623882; DNAm -> cg19045313; DNAm -> cg00719108; DNAm -> cg00517796; DNAm -> cg07537581; DNAm -> cg18426156; DNAm -> cg21202355; DNAm -> cg22839075; DNAm -> cg17961892; DNAm -> cg14019695; DNAm -> cg09126038; DNAm -> cg06529685; DNAm -> cg24080119; DNAm -> cg17847344; DNAm -> cg03978041; DNAm -> cg06514855; DNAm -> cg26383138; DNAm -> cg16877615; DNAm -> cg13126315; DNAm -> cg00730653; DNAm -> cg25623271; DNAm -> cg07366506; DNAm -> cg01173186; DNAm -> cg02225716; DNAm -> cg26404886; DNAm -> cg09002677; DNAm -> cg23601614; DNAm -> cg06684407; DNAm -> cg20586124; DNAm -> cg27102141; DNAm -> cg10805850; DNAm -> cg27364588; DNAm -> cg09602803; DNAm -> cg02319733; DNAm -> cg24702091; DNAm -> cg09914513; DNAm -> cg19727817; DNAm -> cg02498382; DNAm -> cg21624342; DNAm -> cg14866339; DNAm -> cg15866213; DNAm -> cg10736454; DNAm -> cg02661886; DNAm -> cg15646626; DNAm -> cg00081729; DNAm -> cg10333808; DNAm -> cg03070187; DNAm -> cg19783404; DNAm -> cg01382100; DNAm -> cg18504937; DNAm -> cg10427793; DNAm -> cg14366742; DNAm -> cg17437621; DNAm -> cg15336438; DNAm -> cg00047185; DNAm -> cg07003632; DNAm -> cg11942329; DNAm -> cg27357865; DNAm -> cg22429418; DNAm -> cg27085265; DNAm -> cg19701084; (Protein -> NDUFA10_1; Protein -> HSPA8_1; Protein -> DOCK2; Protein -> NDUFV1_2; Protein -> CD44; Protein -> CBX1_2; Protein -> NDUFA5_2; Protein -> SYT7; Protein -> BIN1_4; Protein -> ATP5J2_1; Protein -> PLXNB1; Protein -> PADI2))) | Metabolite -> lysoPC a C20:3; Metabolite -> PC aa C36:6; Metabolite -> LCA; Metabolite -> lysoPC a C17:0; Metabolite -> HDCA; Metabolite -> PC aa C36:5 (RNA -> POLDIP3; RNA -> SARS; RNA -> LOC399715; RNA -> SMTN; RNA -> FRMPD2L2; RNA -> NDUFC2; RNA -> UHRF1; RNA -> FGF18; RNA -> SUNC1; RNA -> HS.213541; RNA -> PRKCH; RNA -> TAS2R10; RNA -> HS.570326; RNA -> DDIT4; RNA -> CLYBL; RNA -> STARD10; RNA -> ADNP2; RNA -> LOC650557; RNA -> MIB2; RNA -> LOC642521; RNA -> PLOD2; RNA -> TUBGCP5; RNA -> FAM59B; RNA -> GAL3ST3; RNA -> EIF2B2; RNA -> PEPD; RNA -> TOR1AIP2; RNA -> CHCHD6; RNA -> ISYNA1; RNA -> C5ORF15; RNA -> HS.583388; RNA -> NOLA2; RNA -> ACOT7; RNA -> TUBA1A; RNA -> GABBR1; RNA -> AQP11; RNA -> HS.145490; RNA -> FGF22; RNA -> SERPINF1; RNA -> ZNF215; RNA -> NAV2; RNA -> NFKB1; RNA -> AFF1; RNA -> AGFG2; RNA -> LOC651403; RNA -> CIB3; RNA -> POLR1E; RNA -> ASCL1; RNA -> NR1I2; RNA -> RFC2; RNA -> CSDE1; RNA -> INO80B; RNA -> PRG-3; RNA -> MED12; RNA -> HS.577965; RNA -> SOX21; RNA -> C11ORF76; RNA -> CDCA5; RNA -> HS.72367; RNA -> NTAN1; RNA -> LOC648364; RNA -> HOXC9; RNA -> UBE2D2; RNA -> LOC647096; RNA -> MPP2; RNA -> HS.567677; RNA -> HS.284464; RNA -> ZIC2; RNA -> HS.218036; RNA -> THOC5; RNA -> MTMR15; RNA -> AK3L1; RNA -> LRP8; RNA -> LOC646424; RNA -> ITPR2; RNA -> HIST1H2AH; RNA -> CLCNKA; RNA -> PRTG; RNA -> PVRL1) DNAm -> cg06481431; DNAm -> cg23449791; DNAm -> cg01553632; DNAm -> cg27609489; DNAm -> cg26366347; DNAm -> cg10678190; DNAm -> cg23044017; DNAm -> cg17949455; DNAm -> cg26361052; DNAm -> cg15830613; DNAm -> cg27059537; DNAm -> cg03383411; DNAm -> cg09298623; DNAm -> cg09798879; DNAm -> cg05505765; DNAm -> cg02081006; DNAm -> cg24433722; DNAm -> cg09329930; DNAm -> cg17512802; DNAm -> cg08439487; DNAm -> cg11348257; DNAm -> cg14757157; DNAm -> cg21463140; DNAm -> cg25932315; DNAm -> cg20777398; DNAm -> cg06145909; DNAm -> cg10269439; DNAm -> cg10679147; DNAm -> cg03139898; DNAm -> cg22682162; DNAm -> cg04515001; DNAm -> cg18050892; DNAm -> cg25691070; DNAm -> cg20638583; DNAm -> cg10215032; DNAm -> cg12652440; DNAm -> cg03907174; DNAm -> cg13602232; DNAm -> cg10409449; DNAm -> cg00934092; DNAm -> cg07470275; DNAm -> cg20659418; DNAm -> cg01021245; DNAm -> cg15927357; DNAm -> cg21438018; DNAm -> cg00425078; DNAm -> cg13704531; DNAm -> cg15899474; DNAm -> cg00056541; DNAm -> cg09114441; DNAm -> cg24675344; DNAm -> cg23288653; DNAm -> cg00500770; DNAm -> cg07604651; DNAm -> cg10607599; DNAm -> cg01527159; DNAm -> cg06636220; DNAm -> cg03713379; DNAm -> cg16514818; DNAm -> cg18225991; DNAm -> cg11946072; DNAm -> cg13657785; DNAm -> cg16588302; DNAm -> cg14186824; DNAm -> cg06177627; DNAm -> cg11361049; DNAm -> cg06238921; DNAm -> cg14531038; DNAm -> cg13715862; DNAm -> cg21772773; DNAm -> cg17918227; DNAm -> cg00711351; DNAm -> cg10128511; DNAm -> cg11491504; DNAm -> cg08108051; DNAm -> cg16293105; DNAm -> cg03339854; DNAm -> cg16465560; DNAm -> cg26672159; DNAm -> cg08459169; DNAm -> cg08573295; DNAm -> cg15643885; DNAm -> cg24319892; DNAm -> cg07850896; DNAm -> cg13162581; DNAm -> cg16622920; DNAm -> cg04144911; DNAm -> cg02919960; DNAm -> cg24673043; DNAm -> cg12695615; DNAm -> cg20644084; DNAm -> cg14872580 Protein -> C9orf16_2; Protein -> BIN1_9; Protein -> BIN1_7; Protein -> PIK3R1_1; Protein -> AP2B1_2; Protein -> SNAP29; Protein -> ELMO1_2; Protein -> MLF2_2; Protein -> SYN1 | Metabolite -> Glu; Metabolite -> Ala; Metabolite -> TwMCA; Metabolite -> Val; Metabolite -> Creatinine; Metabolite -> lysoPC a C RNA -> LOC731950; RNA -> MRPS16; RNA -> RFWD2; RNA -> SNORD56; RNA -> HS.581083; RNA -> HS.583637; RNA -> TBX20; RNA -> HS.156241; RNA -> HS.481464; RNA -> LPIN2; RNA -> UBAP1; RNA -> BRD9; RNA -> LAMA3; RNA -> CLCNKA; RNA -> HS.564742; RNA -> GPKOW; RNA -> HS.145476; RNA -> ARPC5; RNA -> WSB2; RNA -> BCL2L13; RNA -> WBP1; RNA -> HDHD2; RNA -> TOMM40L; RNA -> MANBAL; RNA -> NRSN2; RNA -> GCDH; RNA -> ABCF2; RNA -> DNAJC5B; RNA -> ACSL4; RNA -> C8ORF22; RNA -> HS.545454; RNA -> FIBP; RNA -> PYCRL; RNA -> GSTZ1; RNA -> PARD6G; RNA -> TRIAP1; RNA -> HS.294603; RNA -> JOSD1; RNA -> SDCCAG3; RNA -> CST1; RNA -> SPTBN4; RNA -> BEAN; RNA -> S100A13; RNA -> GABARAP; RNA -> HS.539959 DNAm -> cg06071660; DNAm -> cg19961277; DNAm -> cg27567587; DNAm -> cg16306508; DNAm -> cg21356750; DNAm -> cg14023762; DNAm -> cg05937687; DNAm -> cg03661929; DNAm -> cg03563694; DNAm -> cg14191675; DNAm -> cg10384134; DNAm -> cg04414459; DNAm -> cg12756244; DNAm -> cg09765463; DNAm -> cg19645248; DNAm -> cg05313918; DNAm -> cg21201272; DNAm -> cg03022609; DNAm -> cg03284175; DNAm -> cg00943950; DNAm -> cg04112626; DNAm -> cg08317267; DNAm -> cg22758189; DNAm -> cg16641000; DNAm -> cg19192649; DNAm -> cg08169778; DNAm -> cg20735988; DNAm -> cg10701733; DNAm -> cg07312485; DNAm -> cg15260780; DNAm -> cg06688409; DNAm -> cg09339907; DNAm -> cg20863107; DNAm -> cg12071888; DNAm -> cg19598875; DNAm -> cg19124521; DNAm -> cg07402261; DNAm -> cg06704539; DNAm -> cg00397144; DNAm -> cg12353428; DNAm -> cg21276022; DNAm -> cg08213351; DNAm -> cg14488256; DNAm -> cg07529658; DNAm -> cg05247112; DNAm -> cg23333911; DNAm -> cg00632861; DNAm -> cg24778130; DNAm -> cg10555010; DNAm -> cg00569056; DNAm -> cg03205103; DNAm -> cg13602813; DNAm -> cg25784395; DNAm -> cg21858823; DNAm -> cg24674754; DNAm -> cg12207450; DNAm -> cg27428418; DNAm -> cg00124774; DNAm -> cg24034106; DNAm -> cg10736063; DNAm -> cg15824316; DNAm -> cg19497501; DNAm -> cg26149167; DNAm -> cg14889329; DNAm -> cg26436829; DNAm -> cg14983606; DNAm -> cg07201319; DNAm -> cg21453656; DNAm -> cg25407736; DNAm -> cg16878027; DNAm -> cg04570362; DNAm -> cg03333286; DNAm -> cg20808078; DNAm -> cg07492518; DNAm -> cg02990705; DNAm -> cg17337483; DNAm -> cg24854787; DNAm -> cg27008565; DNAm -> cg15211499; DNAm -> cg24777572; DNAm -> cg25367890; DNAm -> cg26777809; DNAm -> cg22253078; DNAm -> cg14874936; DNAm -> cg02491199; DNAm -> cg03563083; DNAm -> cg07509323; DNAm -> cg24928681; DNAm -> cg09946915; DNAm -> cg05562381 Protein -> CLTA_1; Protein -> LDHA_2; Protein -> BIN1_3; Protein -> STX12; Protein -> STXBP1_3; Protein -> SNAP25_6 |

**eTable 3.** Mean F-values in percentages (95%CIs) for each omic modality by AD subtypes.

| **AD subtype** | **Epigenomic** | **Transcriptomic** | **Proteomic** | **Metabolomic** |
| --- | --- | --- | --- | --- |
|  | **Mean (95%CI)** | **Mean (95%CI)** | **Mean (95%CI)** | **Mean (95%CI)** |
| **1** | 8.8 (8.3-9.4) | 14.1 (12.8-15.5) | 6.9 (5.7-8.0) | 41.5 (35.9-47.1) |
| **2** | 22.4 (20.5-24.3) | 34.6 (31.5–37.6) | 15.1 (11.8-18.3) | 81.2 (69.7-92.6) |
| **3** | 12.3 (11.1-13.4) | 15.5 (14.1-17.1) | 7.1 (5.7-8.2) | 63.7 (39.6-87.8) |

**Note.** Higher f-value show stronger differences from the NCI.

**eTable 4.** Post-hoc comparisons using Bonferroni’s method for inter-subtype differences on the F-value which represents significantly different features from the no cognitive impairment group.

|  | **Subtype-subtype comparisons** | **Difference in means on the F-value (%)** | **95% CI** | | ***p*** |
| --- | --- | --- | --- | --- | --- |
| **Metabolites** | **1 – 2** | **39.7** | **21.6** | **57.7** | **<0.001** |
|  | **1 – 3** | **22.3** | **4.2** | **40.3** | **0.013** |
|  | 2 – 3 | 17.5 | -6.2 | 41.1 | .184 |
| **RNAseq** | **1 – 2** | **20.6** | **17.0** | **24.2** | **<0.001** |
|  | 1 – 3 | 1.6 | 2.6 | 5.7 | 1.00 |
|  | **2 – 3** | **19.0** | **14.9** | **23.2** | **<0.001** |
| **CpGs** | **1 – 2** | **13.9** | **11.9** | **15.8** | **<0.001** |
|  | **1 – 3** | **3.4** | **1.5** | **5.4** | **<0.001** |
|  | **2 – 3** | **10.5** | **8.4** | **12.5** | **<0.001** |
| **Proteins** | **1 – 2** | **8.2** | **5.2** | **11.3** | **<0.001** |
|  | 1 – 3 | 0.09 | -3.4 | 3.6 | 1.00 |
|  | **2 – 3** | **8.1** | **4.4** | **11.8** | **<0.001** |

**Note.** The largest inter-subtype differential F-values relative to NCI were consistently seen between AD subtype 1 and AD subtype 2 with differences in F-values ranging from 8% for proteomic alterations to 40% for metabolomic differences with AD subtype 2 consistently showing higher differential values from AD subtype 1. The smallest inter-subtype differences were observed for AD subtypes 1 and 3 with differences ranging from only 0.1% for proteomics differences to 22% for metabolomic differences differentiating them from NCI. Indeed, no significant differences were observed between AD subtypes 1 and 3 for proportions of RNA and proteomic alterations.

**eTable 5.** Post-hoc comparisons using Bonferroni’s test for mean score subtype-subtype comparisons on neuroticism, depressive symptoms, loneliness, and purpose in life.

| **Measure** | **Subtype-subtype comparisons** | **Difference in means** | **95% CI** | | ***p*** |
| --- | --- | --- | --- | --- | --- |
| **Neuroticism** | **0 – 1** | **2.39** | **0.70** | **4.07** | **0.001** |
|  | 1 – 2 | 1.05 | -1.77 | 2.86 | 0.76 |
|  | 1 – 3 | 0.18 | -1.72 | 2.08 | 1.00 |
|  | 0 – 2 | 1.34 | -3.02 | 0.34 | 0.21 |
|  | 2 – 3 | 0.78 | -2.77 | 1.03 | 1.00 |
|  | **0 – 3** | **2.21** | **0.42** | **4.00** | **0.007** |
| **Depressive symptoms** | 0 – 1 | 0.33 | -0.01 | 0.68 | 0.064 |
|  | 1 – 2 | 0.05 | -0.32 | 0.42 | 1.000 |
|  | 1 – 3 | 0.02 | -0.42 | 0.32 | 1.000 |
|  | **0 – 2** | **0.38** | **0.05** | **0.74** | **0.017** |
|  | 2 – 3 | 0.07 | -0.32 | 0.45 | 1.000 |
|  | 0 – 3 | 0.32 | -0.05 | 0.68 | 0.134 |
| **Loneliness** | 0 – 1 | 0.21 | 0.03 | 0.44 | 0.127 |
|  | 1 – 2 | 0.08 | -0.18 | 0.35 | 1.000 |
|  | 1 – 3 | 0.08 | -0.19 | 0.35 | 1.000 |
|  | 0 – 2 | 0.12 | -0.12 | 0.36 | 1.000 |
|  | 2 – 3 | 0.16 | -0.11 | 0.44 | 0.690 |
|  | **0 – 3** | **0.28** | **0.04** | **0.54** | **0.015** |
| **Purpose in life** | 0 – 1 | 0.17 | -0.01 | 0.34 | 0.073 |
|  | 1 – 2 | 0.01 | -0.18 | 0.21 | 0.21 |
|  | 1 – 3 | 0.01 | -0.19 | 0.21 | 1.00 |
|  | **0 – 2** | **0.18** | **0.001** | **0.35** | **0.048** |
|  | 2 – 3 | 0.02 | -0.18 | 0.23 | 1.000 |
|  | 0 – 3 | 0.15 | -0.03 | 0.34 | 0.171 |

*Note.* 0 denotes the no cognitive impairment group. Bolded rows indicate significant differences.

**Figures**

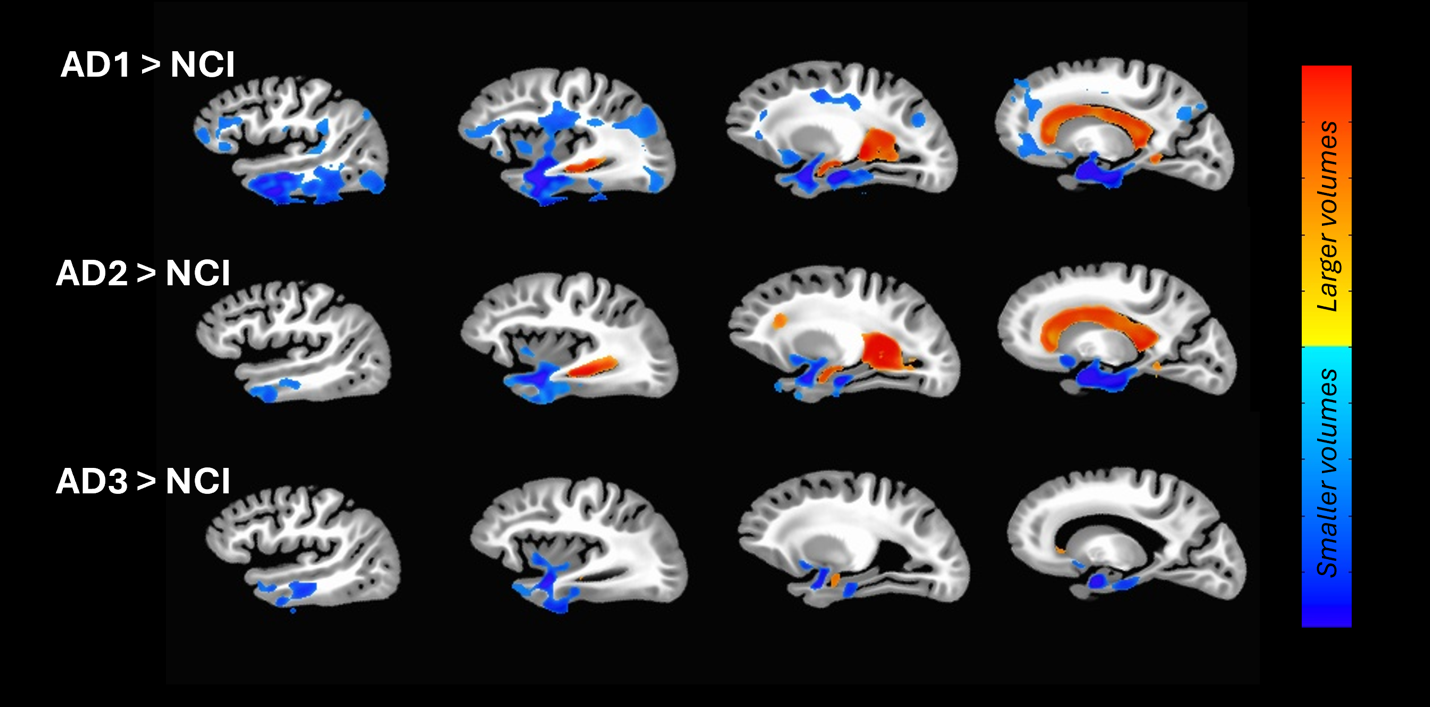

**eFigure 1.** Differences in brain morphology between AD subtypes and the background (FWER *p*≤.05), adjusted for age at death, sex, race, education, postmortem scanning interval and location. Warm and cold colors indicate larger and smaller volumes relative to the background, respectively.
